# Comprehensive functional annotation of metagenomes and microbial genomes using a deep learning-based method

**DOI:** 10.1101/2022.06.06.494653

**Authors:** Mary Maranga, Pawel Szczerbiak, Valentyn Bezshapkin, Vladimir Gligorijevic, Chris Chandler, Richard Bonneau, Ramnik J Xavier, Tomasz Kosciolek, Tommi Vatanen

## Abstract

Comprehensive protein function annotation is essential for understanding microbiome-related disease mechanisms in the host organisms. Still, a large portion of human gut microbial proteins lack functional annotation. Here, we have developed a new metagenome analysis workflow integrating *de novo* genome reconstruction, taxonomic profiling and deep learning-based functional annotations from DeepFRI. We validate DeepFRI functional annotations by comparing them to orthology-based annotations from eggNOG on a set of 1,070 infant metagenome samples from the DIABIMMUNE cohort. Using the workflow, we have generated a sequence catalogue of 1.9 million non-redundant microbial genes. The functional annotations revealed 70% concordance between GO annotations predicted by DeepFRI and eggNOG. However, DeepFRI improved the annotation coverage, with 99% of the gene catalogue obtaining GO molecular function annotations, albeit less specific compared to eggNOG. Additionally, we construct pan-genomes in a reference-free manner using high-quality metagenome assembled genomes (MAGs) and analyse the associated annotations. eggNOG annotated more genes on well-studied organisms such as *Escherichia coli* while DeepFRI was less sensitive to taxa. This workflow will contribute to novel understanding of the functional signature of the human gut microbiome in health and disease as well as guide future metagenomics studies.

## Introduction

The advent of sequencing technologies has resulted in a substantial increase in metagenomic studies sequencing DNA of microbial consortia (microbiomes) inhabiting various hosts and environments. This has led to a significant boost in known microbial genomes contributing substantially to our understanding of the genetic diversity encoded in microbiomes. Although metagenomics provides a great potential to characterize difficult-to-cultivate and uncultivated microbes, our ability to link specific genes to disease phenotypes is hampered by poor understanding of the functions and roles of the majority of the newly discovered microbial genes (Li et al., 2014; Pasolli et al., 2019).

Comprehensive functional annotation is crucial in identifying disease-causing functional changes in proteins, detecting antibiotic resistance genes (Baric et al., 2016; Tuan et al., 2019), and designing new therapeutic strategies. Labor-intensive laboratory experiments still provide the most reliable way of functionally annotating genes and the proteins they encode (Tiwari & Srivastava, 2014). However, due to the rapid increase in the number of uncharacterized proteins, such experimental methods and manual curation fail to scale up to accommodate such a large amount of protein sequence data. This has created a huge sequence-to-function gap which is still widening, because sequencing is high-throughput, while functional characterization continues to be relatively slow. The overall proportion of human gut microbiome protein-coding genes with uncharacterized functions range between 40-60% depending on the annotation method (Li et al 2014; Vatanen et al., 2018). A recent study reported that 27.3% of genes present in the Unified Human Gastrointestinal Genome (UHGG) catalogue could not be mapped to functional databases, while 14.2% of genes matching to COG database were labelled as “function unknown” (Almeida et al., 2021). Automated and scalable methods for microbial protein functions prediction are needed to address this gap.

Most functional annotation methods rely on inferring homology across databases such as Uniprot (Bateman et al., 2021) and the NCBI’s reference sequence (RefSeq) database (O’Leary et al., 2016). Conventional homology-based annotation methods are fast, however, they suffer low functional annotation coverage. Deep learning methods have been considered as an effective complement (Gligorijevic et al., 2019; Kulmanov et al., 2018; Rifaioglu et al., 2019), since they are able to predict protein functions at large-scale irrespective of sequence homology. Metagenome functional annotation can be performed either by assembling sequence reads into contigs, followed by reconstruction of complete and accurate metagenome-assembled genomes (MAGs) and mapping predicted genes to annotated sequences (Quince et al., 2017), or by directly mapping individual reads to annotated gene sequences (Bose et al., 2015; Franzosa et al., 2018; Kim et al., 2016). Various read mapping-based functional annotation workflows exist, including HUMAnN2 (Franzosa et al., 2018), HUMAnN3 (Beghini et al., 2021) and MG-RAST (Keegan et al., 2016). These workflows align short reads directly to a reference catalogue of microbial sequence to estimate functional profiles (Eng et al., 2020). Such approach is, however, limited by the used reference database and fails to annotate genes that lack a homologous counterpart in the reference collection.

Reconstruction of MAGs has proved to be a successful strategy for novel genome discovery and characterization of the functional potential of complex microbial communities (Almeida et al., 2019). The sequencing reads are assembled into long contiguous DNA fragments and further clustered into different bins based on depth of coverage and tetranucleotide frequency across samples (Alneberg et al., 2014; Strous et al., 2012). Several studies have employed this technique providing new insights into the genetic diversity of the human gut microbiome and paving ways for exploring microbial dark matter (Parks et al., 2017; Nayfach et al., 2019).

Widely used schemes for classifying protein functions include the Gene Ontology (GO) (Carbon et al., 2019), Enzyme Commission (EC) numbers (Martínez Cuesta et al., 2015), and Kyoto Encyclopedia of Genes and Genomes (KEGG) (Kanehisa et al., 2017). Gene ontology (GO) terms are attractive as they offer an accurate description of the protein functions levels and relationships between those annotations. The system represents function in a Directed Acyclic Graph (DAG) type relationship where protein attributes are divided into three main categories: molecular function (MF), biological process (BP) and cellular component (CC). Importantly, GO terms facilitate more comprehensive inter-method comparisons through these protein annotation relationships even when the specificity of annotations differ between methods.

Here we describe a new metagenome assembly workflow integrating *de novo* genome reconstruction, taxonomic profiling and deep learning-based gene annotations from DeepFRI (Gligorijevic et al., 2021). We validate the functional annotations by comparing them to orthology based annotations from eggNOG (Huerta-Cepas et al., 2019). We further show that such an approach almost exhaustively annotates millions of metagenome-assembled genes, although the annotations are, on average, less specific compared to the homology-based annotations from eggNOG. Finally, we demonstrate integration of gene annotations and metagenome-assembled pangenomes using >1,000 infant metagenomes from the DIABIMMUNE cohort. In summary, combining gene annotations from DeepFRI with the commonly used orthology-based annotations helps understanding the roles of novel microbial genes observed in metagenomic surveys and could eventually close the sequence-to-function gap hindering most microbiome studies.

## Results

### Workflow architecture

The workflow is fully automated for metagenomic assembly, binning of metagenome-assembled genomes, construction of gene catalogue and functional annotation. It integrates state-of-the-art bioinformatic tools via Docker containers. The workflow is implemented using Workflow Description Language (WDL) and allows flexibility of bioinformatic tools versioning and scalability of memory in high-performance computing environments. This allows large-scale functional annotations of metagenomics data by leveraging high-quality protein information to annotate functions with higher coverage. It takes raw paired-end Illumina reads (short reads) as input and performs data analysis in five phases (1) Assembly of sequencing reads into contigs, gene prediction and clustering to generate a gene catalogue, (2) functional annotation of the gene catalogue, (3) Binning of assembled contigs into metagenome-assembled genomes (MAGs), (4) Taxonomic annotation of MAGs and (5) Mapping between MAGs and functionally annotated gene catalogue (Figure 1). We have developed a custom Python script for mapping between metagenomic species and functionally annotated gene catalogue. On completion, the workflow provides various tabular output files such as the metagenome species (MAGs) constructed, non-redundant gene catalogue, functional annotations and gene mapper table as tab-separated output files.

**Figure 1:**
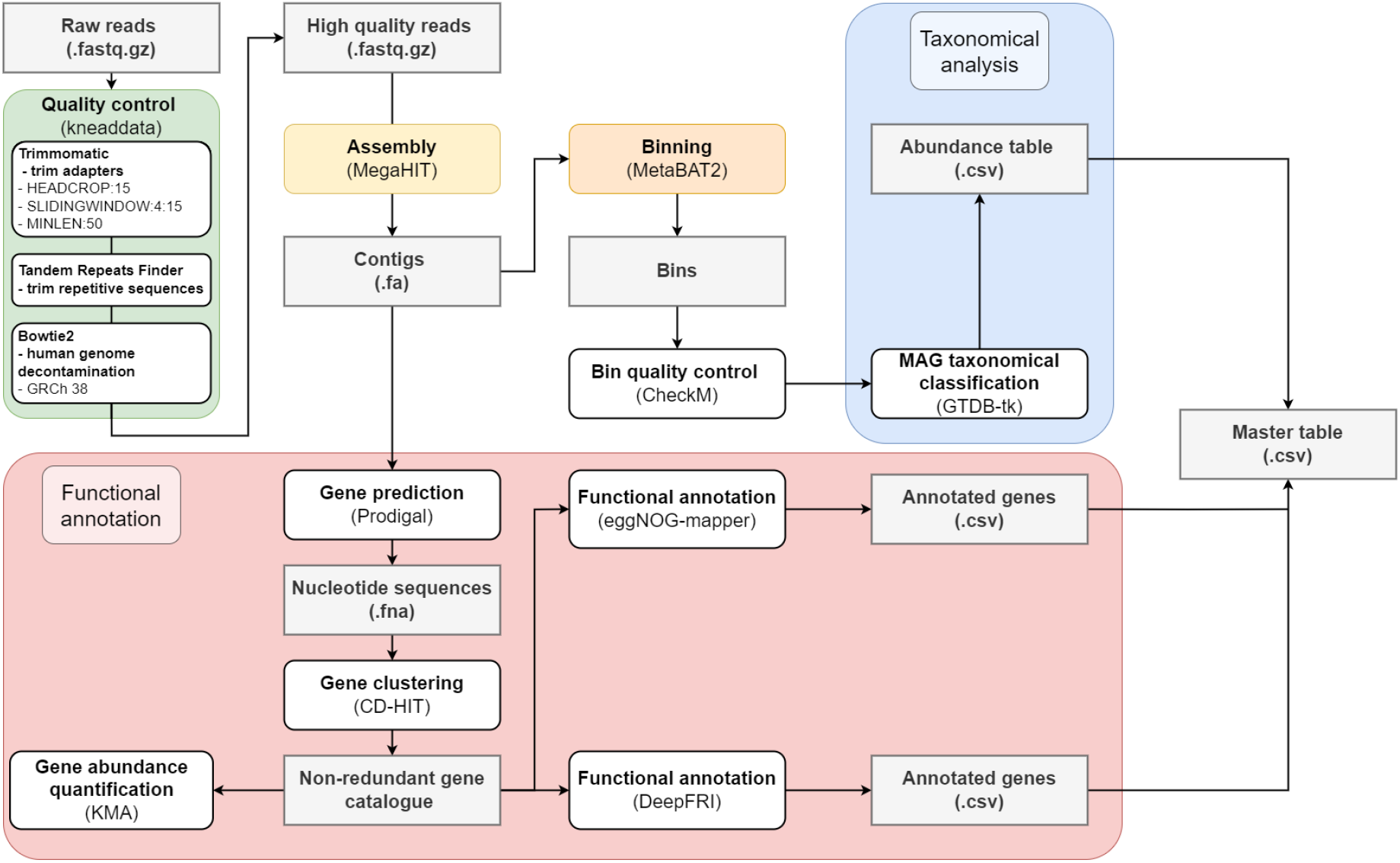
Schematic workflow overview.

### Undescribed diversity in infant gut metagenomes

To demonstrate feasibility and robustness and to highlight several use cases of the workflow data products, we analyzed metagenomic data from the DIABIMMUNE cohort (Vatanen et al., 2019) (https://diabimmune.broadinstitute.org/diabimmune/). This constituted a longitudinal sample series from individual infants and young children from Finland, Estonia and Russia. Overall, we aggregated metagenome shotgun sequencing data from 1,070 samples including 202 samples from Estonia, 586 samples from Finland and 282 samples from Russia. Average number of reads per sample was 1.49 × 10^7^ paired end reads (minimum 9.28 × 10^3^; maximum 8.09 × 10^7^).

The first step for functional analysis was the construction of a non-redundant gene catalogue. We performed *de novo* assembly of the metagenomes resulting in a total of 17 million long contigs of length ≥500 bp mean 16,510 contigs per sample (minimum 94; maximum 60,979 contigs), collectively harboring approximately 21.9 million ORFs, as predicted by Prodigal. Clustering of these genes into gene families with >95% sequence similarity resulted in a catalogue of 1.9 million non-redundant genes (Table 1).

**Table 1:**
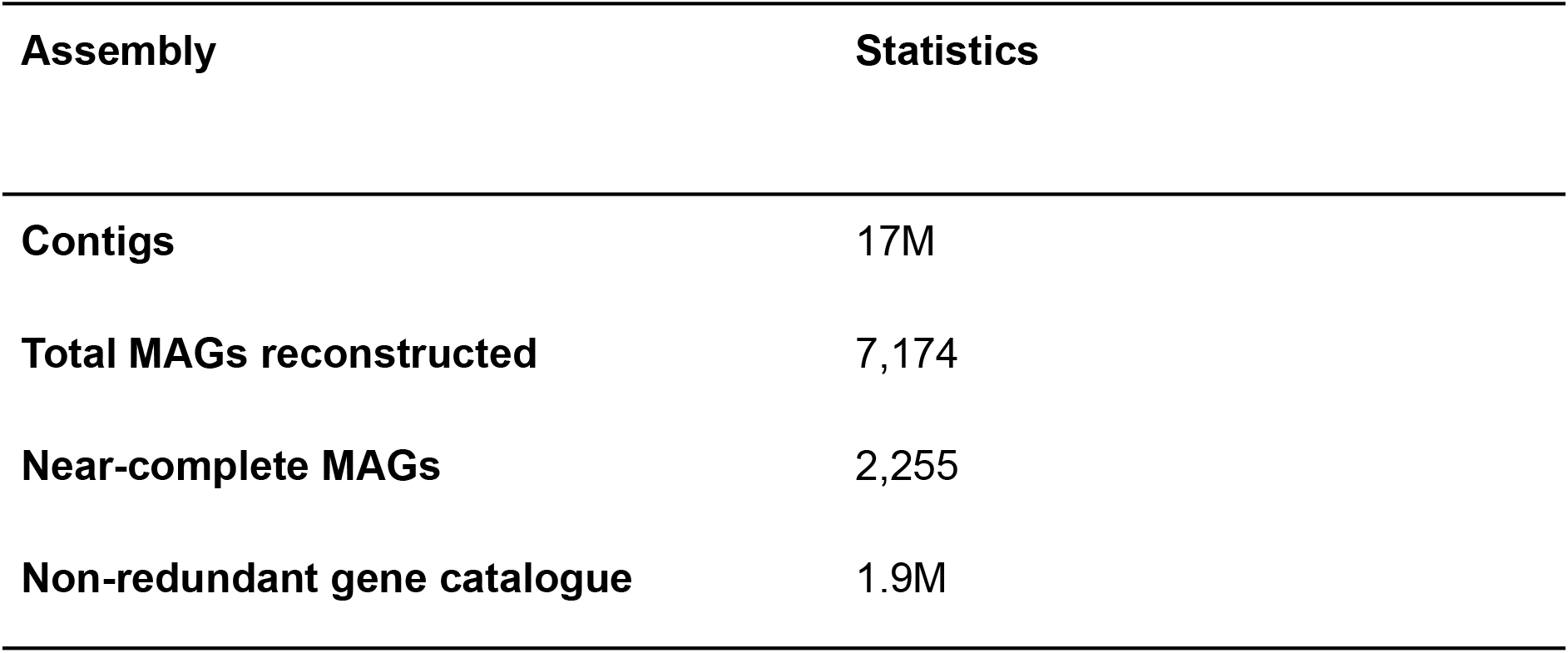
Assembly and gene prediction statistics for the Diabimmune dataset.

| Assembly | Statistics |
| --- | --- |
| Contigs | 17M |
| Total MAGs reconstructed | 7,174 |
| Near-complete MAGs | 2,255 |
| Non-redundant gene catalogue | 1.9M |

Metabat binning of contigs produced a collection of 7,174 MAGs, of which 2,255 were near-complete bacterial genomes (>90% completeness, <5% contamination) (Table 1 and Supplementary figure 1) which corresponded to 2 near-complete MAGs per sample, on average. Taxonomic annotation of each genome was carried out using GTDB-tk. The high-quality genomes spanned 202 bacterial species, with most MAGs representing phyla *Firmicutes* (884 genomes) *Bacteroidota* (823 genomes), *Actinobacteriota* (296) and genera *Bacteroides* (614 genomes), *Bifidobacterium* (268 genomes), and *Faecalibacterium* (124 genomes) (Supplementary figure 2b-c).

To visualize the distribution of high quality near-complete genomes across phylogeny, we build a maximum likelihood phylogenetic tree based on high quality near-complete genomes. The tree was constructed by PhyloPhLAN (Asnicar et al., 2020) and visualized using Interactive Tree of Life (v5.6.2) (Letunic & Bork, 2019). The genomes spanned 10 bacterial class, including *Coriobacteriia, Actinomycetia, Bacilli, Negativicutes, Clostridia, Campylobacteria, Alphaproteobacteria, Gammaproteobacteria, Verrucomicrobiae*, and *Bacteroidia* (Figure 5a). We observed high phylogenetic dispersion in *Faecalibacterium* genera. *Faecalibacterium* has been shown to be highly diverse and comprise multiple different phylotypes (Pasolli et al., 2019).

To integrate information on gene homology from the gene catalogue and taxonomic information obtained through MAGs, we tried to provide further insights into the following question: are 95% gene families taxon specific or shared across genomes of different species, genera or even families?. Interestingly we found that a notable portion of the gene families had multiple taxonomies assigned, 55,006 (3%) at family level, 208,885 (11%) at genus level, and 276,042 (14%) at species level (Supplementary figure 2a). We hypothesize that shared genes could be attributed to the horizontal gene transfer between species although additional studies are required to conclusively confirm this.

### Functional representation of the DIABIMMUNE infant metagenomes

To elucidate the functional composition of the gut microbiota we annotated genes using DeepFRI and eggNOG. We focussed on the molecular function branch of Gene Ontology and only considered DeepFRI annotations with a probability threshold above 0.2 to be significant. We first validated the predicted annotations by comparing the level of concordance between both methods based on the specificity of the GO terms, expressed as Shannon information content. The information content (IC) is used to quantify the specificity of a GO term in the context of the entire set of annotations, such that GO terms annotating many genes are considered to be general and hence contain low information content while rare occurring GO terms are considered more specific and contain higher information (IC values) (Gaudet et al., 2009). We propagated the Gene Ontology annotations upward through the child-parent relationships using a list of information content values obtained from (Gligorijevic et al., 2021), where IC was calculated as follows:

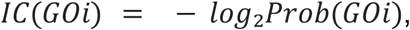

where, Prob(GO_*i*_) is the probability of GO term *i* occurring in the UniProt-GOA database (*n*_*i*_/*n*, where *n*_*i*_ equals number of annotations with a term *i* and *n* is total number of annotated proteins in UniProt-GOA) (Gligorijevic et al., 2021).

By comparing consensus in the gene ontology annotations predicted by both methods, we observed a gradual decrease in the number of genes with high information content annotation in DeepFRI compared to eggNOG (Figure 2a). DeepFRI gave a high number of general annotations while eggNOG was shown to predict more specific MF-GO terms (IC > 12). Next, we wanted to assess whether genes annotated by both methods gained function at the same information content level. We further stratified annotations into different categories; (1) concordance annotations, where eggNOG and DeepFRI annotations agree (genes obtained functions at the same information content level); (2) discordant, where eggNOG and DeepFRI annotations disagree but annotations from both methods exist; (3) DeepFRI and (4) eggNOG unique, genes obtained function from only one method. Our results demonstrated high concordance annotations between eggNOG and DeepFRI (Figure 2b-c). Of all the genes annotated by both methods, the proportion of concordance annotations was 70%, on average. Additionally, DeepFRI had a high number of unique annotations thus expanding the functional landscape of metagenome-assembled genes.

**Figure 2:**
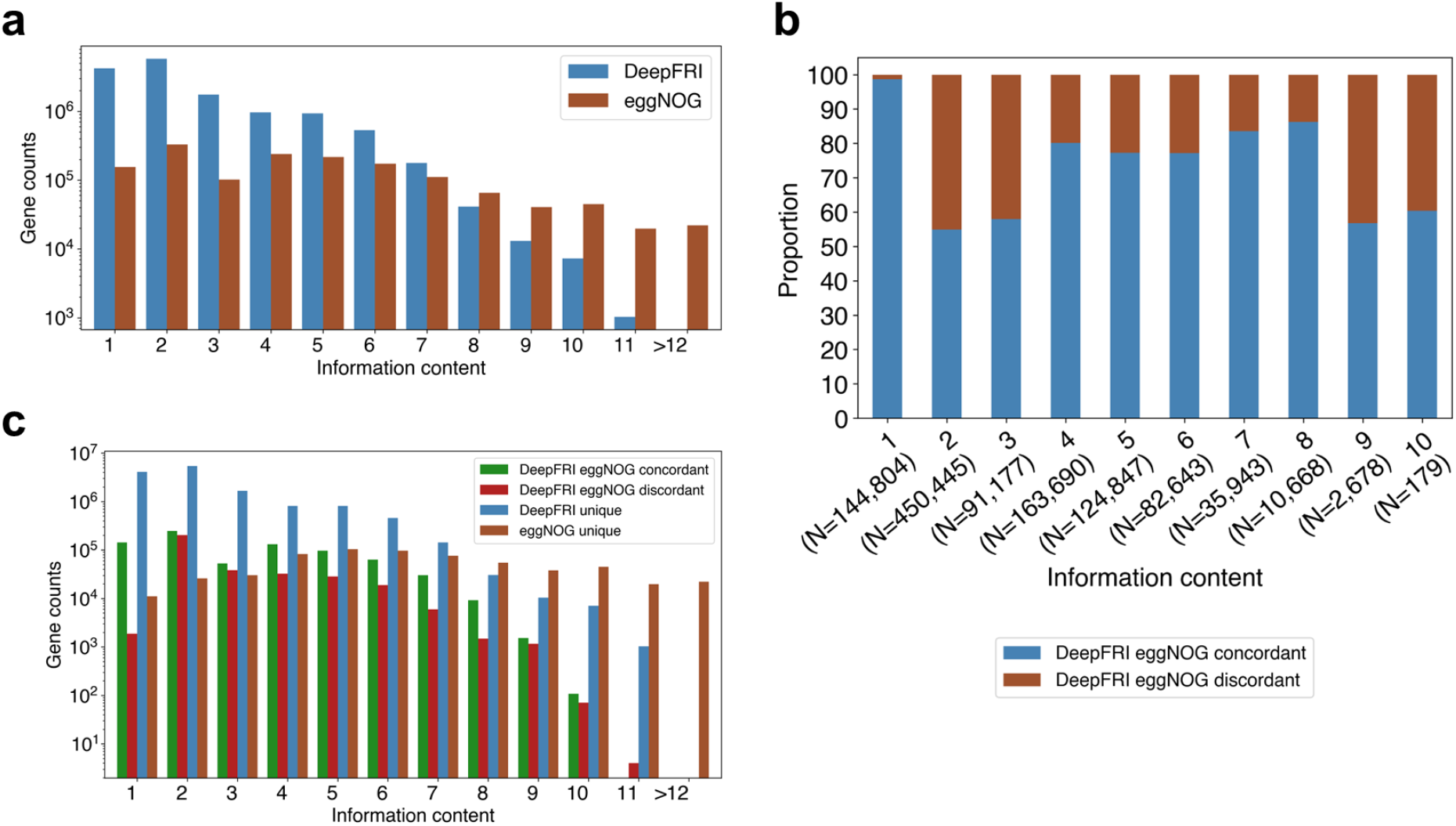
Concordance between DeepFRI and eggNOG annotations. **(a)** Difference in the annotation information content between eggNOG and DeepFRI. (**b)** Proportion of concordant and discordant annotations between DeepFRI and eggNOG per each information content level. (**c)** Consensus between DeepFRI and eggNOG annotations. See Table S3 for the list of information content values.

We then looked into functions encoded in the microbial genes. In total nearly all the genes in the gene catalogue (1,895,540; 99%) obtained GO molecular function annotations by DeepFRI at 0.2 probability threshold whereas only (219,634; 12%) of genes obtained similar annotations by eggNOG (Table 2). Further, we complemented the annotations by considering functional annotations from eggNOG free text description which consolidates information obtained from several source databases such as SMART/Pfam (Huerta-Cepas et al., 2019). A total of 1,372,653 (72%) genes obtained functional information from eggNOG free text description. We then compared differences in gene sets annotated by both methods. The Venn diagram (Figure 3b) shows the intersection among genes annotated by DeepFRI and eggNOG, with 219,003 genes in common across the two gene sets. A total of 1.6M genes were unique to DeepFRI and 631 genes unique to eggNOG gene ontology. Additionally, 1.3M gene sets were common across DeepFRI and eggNOG free text description annotations (Figure 3c).

**Table 2:** Genes annotated by DeepFRI and eggNOG.

| Method | DeepFRI<br>molecular<br>function | eggNOG<br>Molecular<br>function | eggNOG<br>free text description |
| --- | --- | --- | --- |
| Annotated<br>genes | 1,895,540 (99%) | 219,634 (12%) | 1,372,653 (72%) |

**Figure 3:**
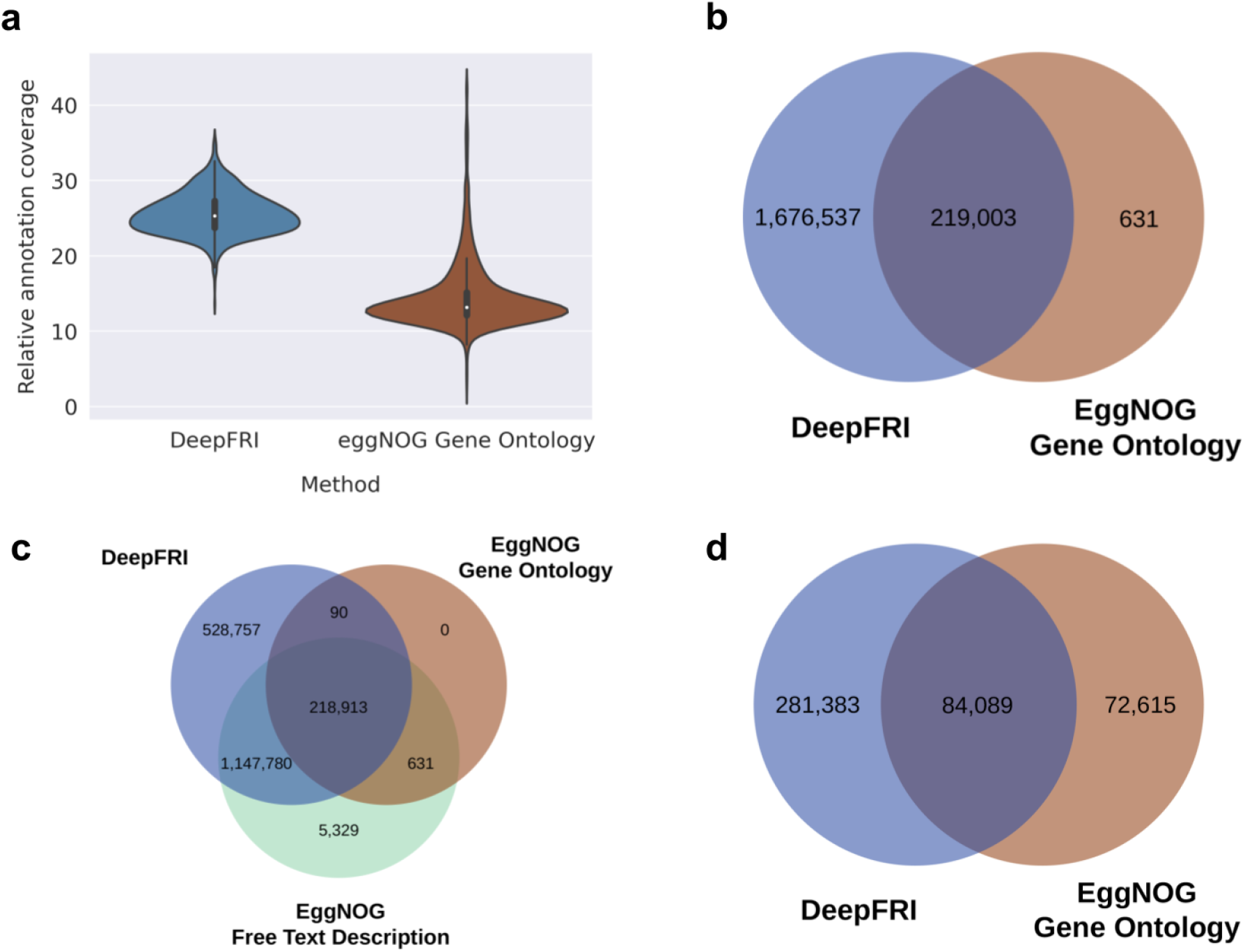
Functional capacity of the gut microbiota. **(a)** Proportion of metagenomics gene abundance with functional annotation by DeepFRI and eggNOG (informative gene ontology terms). The annotation is weighted by relative abundances normalized in copies per million. (**b)** Venn diagram of the number of gene sets annotated by DeepFRI and eggNOG gene ontology (all GO terms). (**c)** 3-way comparisons of gene sets annotated by DeepFRI, eggNOG gene ontology (all GO terms) and eggNOG free text description. (**d)** Venn diagram comparisons of gene sets annotated by DeepFRI and eggNOG using only informative gene ontology terms.

The gene ontology protein function description is represented as a directed acyclic graph (DAG), such that a gene/protein can have multiple GO terms annotated, and a term can have multiple relationships to broader parent terms and more specific child term (child-parent relationship). For example, the molecular function term ‘ATP binding’ has parent terms ‘binding’. For downstream analysis, we filtered out the general gene ontology terms using a subset of 614 informative GO term list obtained from previous work by Vatanen et al., 2017, following the method proposed in (Huang et al., 2007; Zhou et al., 2002). The informative GO categories are obtained by traversing the GO tree from the root and selecting terms that satisfy the following parameters; (i) terms that are associated with more than k=2,000 proteins and (ii) each of their descendant term contains less than 2,000 proteins (k=2,000 equates to approximately 1 of every 5,000 UniRef50 protein families). This gene ontology informative set provides more resolution to extensively studied processes. Filtering of gene ontology terms resulted in 365,472 genes annotated by DeepFRI and 156,704 genes annotated by eggNOG. The Venn diagram (Figure 3d) shows the intersection of 84,089 genes common between DeepFRI and eggNOG gene ontology. A total of 281,383 genes were unique to DeepFRI and 72,615 genes unique to eggNOG.

Finally, we evaluated the relative abundance of annotated genes between DeepFRI and eggNOG considering only genes annotated with informative gene ontology sets. We computed the proportion of metagenomic gene abundance with functional annotation as follows: (1) paired-end reads are mapped to the gene catalogue using the KMA tool, (2) per-gene alignment statistics are then weighted based on the target sequence length to produce abundance values for each gene family. We observed an increase in the annotation coverage at metagenome level by DeepFRI with an average coverage of 25.6% in comparison to eggNOG which had an average coverage of 14.5% (Figure 3a). The main molecular functions annotated by DeepFRI and eggNOG are ATP binding (GO:0005524), structural constituent of ribosome (GO:0003735) and zinc ion binding (GO:0008270) (Supplementary figure 4).

### Pan-genome diversity patterns within the infant metagenomes

The pan-genome can be defined as the entire gene repertoire of all strains in a species (Tettelin et al., 2005). Genes in a pan-genome are classified into two categories: core and accessory genes. The core genes are shared by genomes within a species while accessory genes are present in a subset of the genomes within a species. The pan-genomes were constructed in the following manner: we recruited near-complete metagenome assembled genomes (≥90% completeness and <5% contamination) and included species with at least ten independent near-complete MAGs. We then considered the genes present in ≥90% of MAGs of each species as “core” genes (accounting for the incompleteness of the MAGs), while genes present in <90% of MAGS were considered as “accessory” genes. Combined core and accessory genes constituted the pan-genome of a species.

The pan-genomes covered a total of 42 bacterial species (Supplementary figure 3a) and consisted of a total of 70,997 core genes (on average, 1,690 core genes per pan-genome, ranging from 573 to 3,612 genes) and 355,761 accessory genes (on average, 8,470 accessory genes per pan-genome, ranging from 1,222 to 27,879 genes) (Supplementary figure 3b and Supplementary table 1). Each species had distinctly different core genome and pan-genome sizes. Our pan-genome analysis showed that *Veillonella parvula* and *Bifidobacterium pseudocatenulatum* contained the smallest core genomes with 573 and 753 genes respectively. In contrast, *Escherichia coli* contained the largest core genome (3,612 genes). To investigate differences in intra-species gene richness, we analysed the pan-genome size in relation to the number of genomes (MAGs). We found a strong correlation between the number of MAGs and the size of the pan-genome, with a Pearson correlation r=0.8 (Figure 4c). *Bacteroides ovatus* and *Blautia wexlerae* displayed a larger pan-genome size than expected by the trend. Bacteroides species have been shown to have large pan-genome sizes (Zou et al., 2019). Their remarkable genome plasticity facilitates adaptation to various ecological niches, and interaction with the host immune system (Bencivenga-Barry et al., 2020). *Akkermansia muciniphila, Bifidobacterium longum* and *Bifidobacterium bifidum* had a smaller pan-genome than expected by the trend.

**Figure 4:**
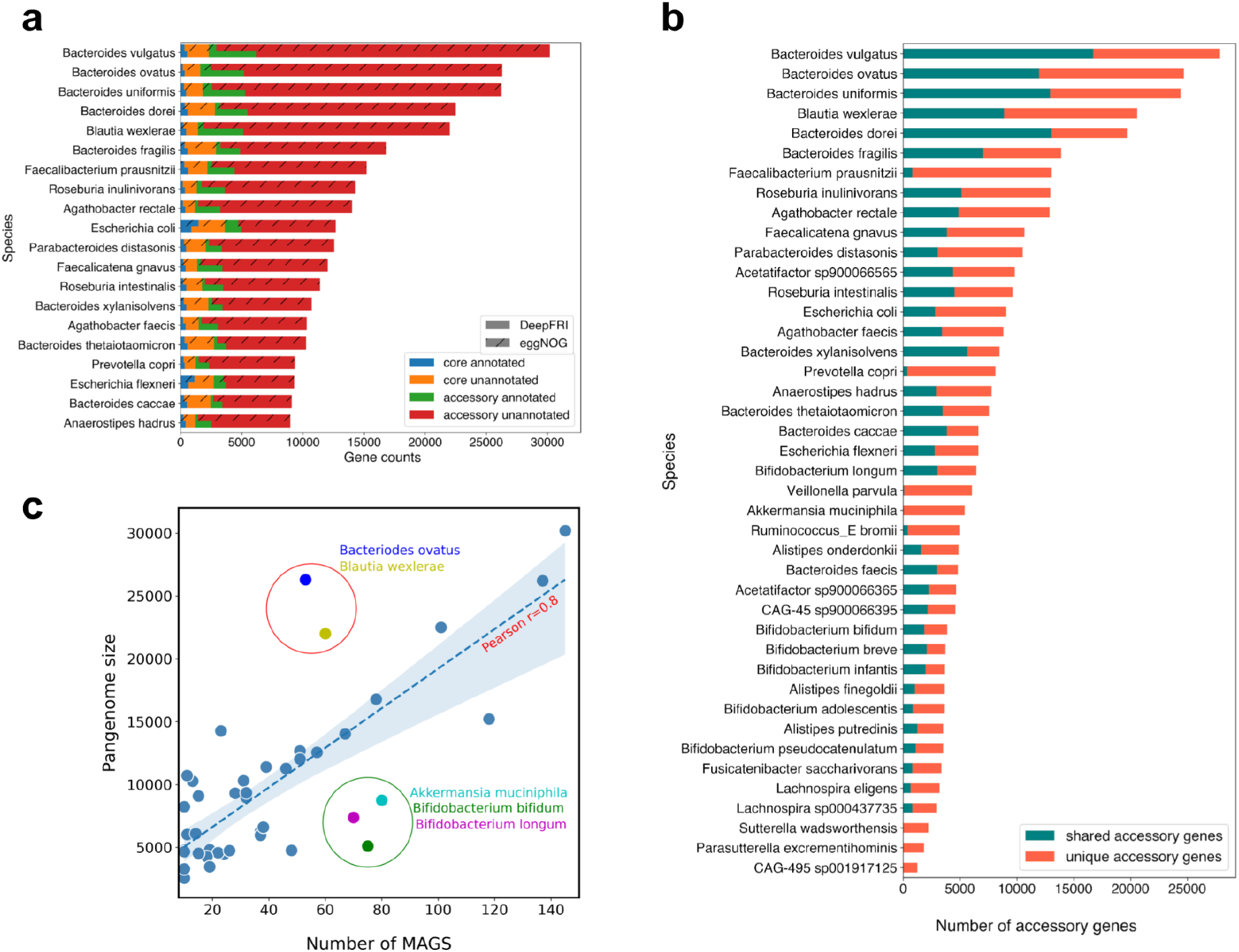
Pan-genome patterns within the infant metagenomes. (**a)** Size of the core and accessory genomes per species stratified by the functional annotation of genes using eggNOG and DeepFRI (known versus unknown function). Entries are ordered according to the size of the pangenome (20 out of 42 species used to construct the pangenomes). (**b)** Number of unique (not shared with other species) and shared accessory genes per pangenome. See Table S1 for a full pan-genome size list. (**c)** Pan-genome size in relation to the number of genomes (MAGs)

To explore the diversity within the pangenomes, we compared the accessory gene repertoire of each species. We observed a wide variation in the gene content between various species. *Bacteroides* species had the highest number of shared accessory genes (accessory genes that were observed in more than one species, Figure 4b). This observation is consistent with a recent study that showed that *Bacteroides* species exchange genes within the genus more frequently through horizontal gene transfer compared to other species such as *Bifidobacterium* (Groussin et al., 2021). Interestingly, species such as *Veillonella parvula, Akkermansia muciniphila, Parasutterella excrementhominis*, and *Sutterella wadsworthensis* accessory genomes harboured very low numbers of shared genes; 76, 56, 6 and 1, respectively, and the rest of the accessory genome was unique (Figure 4b and Supplementary table 1). *Bacteroides dorei* is one of the most diverse and prominent members in the infant gut playing an essential role in immune activation (Vatanen et al., 2017, Vatanen et al., 2019). We wanted to look into its genomic diversity by comparing its accessory genome to other *Bacteroides* species. The analysis showed that *Bacteroides dorei* harboured 7,514 unique accessory genes (accessory genes not found in any other species) representing 38% of its accessory genome (Supplementary figure 3c). One plausible explanation for this diversity could be caused by gene gain through horizontal gene transfer.

To obtain a better understanding of the functions encoded in the core and accessory genomes, we annotated the pan-genomes using DeepFRI and eggNOG. Here, we used only informative GO terms. There was a noticeable difference between annotations predicted by DeepFRI and eggNOG. Our results show that eggNOG annotated more functions on well-studied genomes *such as Escherichia coli*. The core genome of *Escherichia coli* had the highest annotation level, (1,449; 40%) of core genes obtaining GO term annotations from eggNOG and (880; 24%) from DeepFRI (Figure 4a). This could be attributed to the fact that *E. coli* is an extensively studied model organism (Ghatak et al., 2019). Moreover, this observation was supported by our phylogenetic analysis that showed that eggNOG annotated more genes belonging to *Gammaproteobacteria* (Figure 5b). On the other hand, DeepFRI annotations were less sensitive to taxa although DeepFRI annotated more genes belonging to members of *Bacteroides* genera. *Bacteroides vulgatus* species had the highest number of annotated genes (4,383;7%) compared to eggNOG which annotated (928;4.5%) genes (Figure 4a and Figure 5a). Our analysis provides detailed identification of the functional profile of these species and can guide future studies aimed at uncovering potential mechanisms responsible for the diversity in *Bacteroides*.

**Figure 5:**
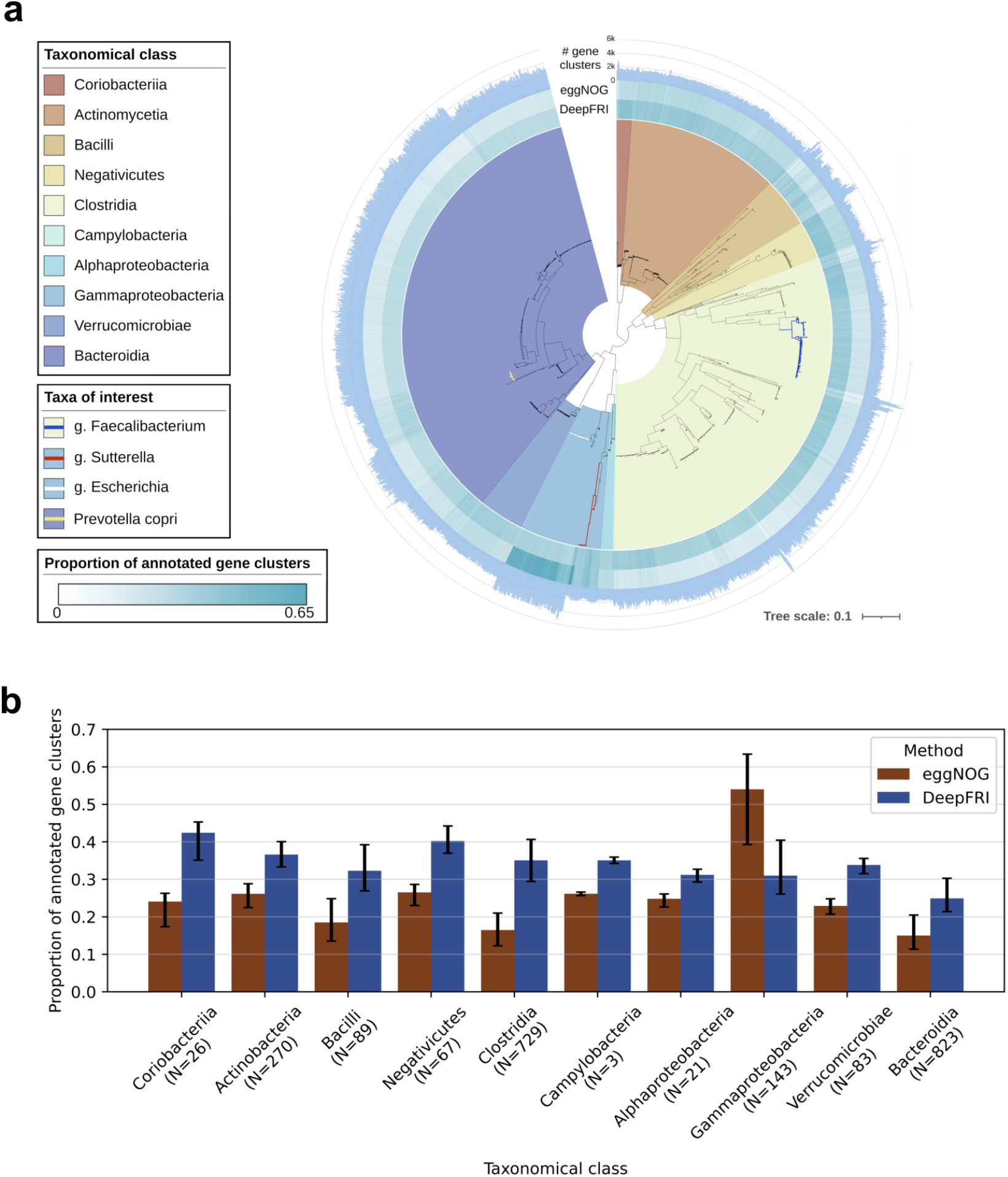
Phylogenetic analysis and taxonomic annotation of high-quality metagenome-assembled genomes. **(a)** Maximum-likelihood phylogenetic tree of 2,255 high-quality near-complete genomes. The taxonomy of the MAGs was assigned by GTDB-Tk. The innermost layer corresponds to 10 bacteria class. The second and third ring represent the proportion of genes annotated by DeepFRI and eggNOG. Bars in the outermost layer indicate the number of gene families per MAG. (**b)** Distribution of annotated genes per class.

### Antibiotics-related GO terms

Finally, we investigated the antibiotics specific biological process GO terms. DeepFRI annotated only two processes, antibiotic catabolic process (GO:0017001) and antibiotic metabolic process (GO:0016999). The antibiotic catabolic process was identified in 32 genes belonging to 22 species, while antibiotic metabolic process was observed in 12 genes coming from 12 species (Supplementary table 4b). Contrastingly, EggNOG annotated more antibiotics-specific GO terms, such as bacteriocin metabolic process (GO:0046224), beta-lactam antibiotic catabolic process(GO:0030655), beta-lactam antibiotic metabolic process (GO:0030653), cephalosporin metabolic process (GO:0043645), macrolide metabolic process (GO:0033067) and response to antibiotics (GO:0046677) (Supplementary table 4a).

Bacteria secrete bacteriocin peptides giving them a distinct advantage to compete against other bacteria present in the same ecological niche. These peptides are known to have a rapid antimicrobial activity towards closely related strains and multi-drug resistant pathogens (Cotter et al., 2013; Jiang et al., 2017). We identified the bacteriocin metabolic process in *Bacteroides, Lactobacillus plantarum* and *Sutterella wadsworthensis*. Bacteriocins produced by *lactobacillus plantarum* have been reported to display antimicrobial activity against both Gram-positive bacteria and Gram-negative bacteria, *Staphylococcus aureus, Escherichia coli, Pseudomonas aeruginosa* (Kumar et al., 2016)

*Bacteroides* species have a high genome plasticity and show increasing resistance rates to multiple antibiotics, including beta-lactams and tetracycline (Rong et al., 2021; Veloo et al., 2019). Our results show that *Bacteroides* possess multiple genes involved in response to antibiotic (GO:0046677), macrolide biosynthetic process (GO:0033068) and cellular response to antibiotic (GO:0071236) 163, 12, 27 genes respectively. *Bacteroides* is a biologically diverse clade and further studies in antibiotic resistance adaptations in *Bacteroides* species are warranted.

## Discussion

Accurate functional assignment of metagenomic sequences is crucial for unlocking the microbiome clinical potential. Here we provide a standalone and fully automated workflow implementing metagenomic assembly, construction of MAGs and non-redundant gene catalogue and comprehensive gene function predictions. The workflow performs different steps of metagenomics analysis in a highly reproducible and scalable environment using Workflow Description Language (WDL). Its features provide functional inference of each gene, including taxonomic classification through better linkage of marker genes in the GTDB reference tree, as well as visualization of near-complete genomes. This allows robust characterization of the functional potential encoded in the microbial genomes.

To demonstrate the usage of our workflow, we tested it on infant gut metagenomes from the DIABIMMUNE cohort (Vatanen et al., 2019). Metagenomic single-sample assembly strategy employed here (Li et al., 2015), represents a scalable methodology for large-scale metagenomes analysis and profiling of functional diversity within the microbial communities. Additionally, the workflow conducts binning of contigs to high quality genomes using Metabat2 (Kang et al., 2019). We recovered MAGs with high completeness (≥90%) and low contamination (<5%). Construction of complete or near-complete MAGs have enabled identification of valuable new genomes from rare species and quantification of intrapopulation diversity (Albertsen et al., 2013; Bowers et al., 2017; Wibowo et al., 2021). It has also been shown to identify novel genes with distinct functional properties associated with human diseases (Nayfach et al., 2019).

For functional annotation we utilized the advantage of deep learning method ability to process massive amounts of data. Graph convolutional networks (GCN)-based DeepFRI ensures efficient and accurate mapping of protein sequences to function annotation at a large scale. It enables discovery of novel protein functions by combining two aspects of information, features learned from protein sequences and contact graphs derived from 3-dimensional structures (Gligorijevic et al., 2021). Combining DeepFRI with commonly used orthology-based method eggNOG not only improves our annotation capabilities but also helps us understand function better. Moreover, we provide an additional layer of function information obtained from eggNOG such as Smart/Pfam annotations (Mistry et al., 2021).

The functional diversity within the human gut microbiome remains poorly characterized as most genes still lack functional annotation (Almeida et al., 2021). By leveraging high quality protein information, we demonstrate that DeepFRI significantly improved the functional annotation coverage of the assembled genes (99%). Further validation of the GO terms predicted by DeepFRI showed a high level of concordance with eggNOG annotations, 70% on average. This indicates that DeepFRI produces reproducible and relevant predictions for the biological interpretation. We show that despite DeepFRI giving less specific (generic) GO term annotations, it provides a more comprehensive functional landscape of almost all metagenome-assembled genes. This increase in annotation coverage represents a key step towards novel understanding of the human gut microbiome, thus alleviating the existing sequence-to-function knowledge gap. Moreover, we provide an important functional resource for future interventions leveraging the gut microbiome to improve health.

Another critical aspect of our workflow is the full support of pan-genome analysis. Pan-genome analysis provides an undiscovered wealth of information for studying the diversity within species. Here we provide a way to construct pangenomes in a reference-free manner from near-complete metagenome assembled species (≥90% completeness and <5% contamination). MAGs serve as an important resource in pan-genome analyses, especially where unculturable species or species without reference genomes are being studied. The constructed pan-genomes were of high quality and thus provided a better understanding of the functional repertoire encoded in core and accessory genes. The functional predictions generated from the species pan-genomes revealed a striking difference between DeepFRI and eggNOG annotations. Notably, DeepFRI annotated more functions coming from members of *Bacteroides* species. The pangenome analysis done here could be leveraged for further studies to understand species-specific gene functions or identification of acquired functions of biomedical relevance such as antibiotics resistance.

There are certain limitations of the workflow. First, we only tested our workflow on the infant gut metagenomes. More robust evaluation is required to demonstrate the performance on other complex metagenomes such as human skin metagenomes, environmental microbiome (seawater/soil metagenomes) and other specific communities. Second, DeepFRI only provides Gene Ontology annotations, therefore, it is difficult to profile antibiotic resistomes (antibiotic-resistant genes), as well as genes encoding carbohydrate-active enzymes (CAZymes), which are important in health and disease.

Given the potential and limitations highlighted above, future integrations of high-quality structure information coming from DeepFRI GCN predictions will allow large-scale functional annotation of metagenomics data with higher accuracy. Additionally, we plan to incorporate KEGG orthology (Kanehisa et al., 2017) and evaluate genes and pathways associated with disease.

We show that the workflow is robust and reproducible for large-scale metagenomics functional annotation, where short read sequences can be fully processed into annotated files for downstream analysis and visualization. The workflow is available as a Docker image and makes use of standard tools for metagenome analysis and functional annotation. Implementation of the workflow in WDL allows extensive parallelization using high performance computing and cloud computing environments. Additionally, its modular architecture set-up enables tool versioning while allowing users to tweak different parameters to suit their specific needs. The Docker images used in the workflow are publicly available at https://github.com/microbiome-immunity-project/metagenome_assembly.

## Methods

### Quality control and assembly

Metagenomics data usually contains sequencing artifacts such as low-quality reads, host contamination and adapter sequences, which compromise downstream analysis. The quality control phase utilizes kneadData https://huttenhower.sph.harvard.edu/kneaddata/. KneadData combines Trimmomatic for adapter sequence and low-quality base trimming (Bolger et al., 2014), Bowtie2 (Langmead & Salzberg, 2012) for removal of host-derived reads and Tandem Repeat Finder for removing tandem repeats in DNA (Benson, 1999).

After the quality control process, the workflow uses MegaHIT (v2.4.2) (Li et al., 2015) to assemble each sample individually into contigs. MegaHIT has been shown to reconstruct known genomes accurately by employing data structures known as the *de Bruijn* graph which decomposes reads into k-mers, thus using minimal memory requirements (Quince et al., 2017; Sczyrba et al., 2017). Additionally, it captures more genes allowing higher resolution of functional profiling and phenotype to genotype analysis of specific microbial communities. The assembled contigs less than 500 bp long are filtered out to yield high-quality contigs.

### Construction of the gene catalogue and gene abundance estimation

For gene annotation, open reading frames from each contig (length ≥ 500 bp) are predicted using Prodigal (Hyatt et al., 2010). The gene products are then clustered at 95% sequence similarity using CD-HIT (v4.8.1) (Fu et al., 2012) to generate the non-redundant gene catalogue. Thereafter, gene abundance quantification is computed by mapping paired-end reads from each sample against the gene catalogue using KMA tool (Clausen et al., 2018). Hits to a sequence are weighted based on the target sequence length and further normalized to produce counts in copies per million (CPM) units.

### Binning of contigs to metagenome-assembled genomes and taxonomic classification

Construction of metagenome assembled genomes provides novel biological insights into genetic diversity within the microbial communities. Our workflow implements an adaptive binning approach using Metabat2 (v.2.3.0) (Kang et al., 2019) by clustering contigs into bins (metagenome-assembled genomes, MAGs) based on the use of tetra-nucleotide frequencies, differential abundance and coverage across samples. The workflow supports binning for each sample individually thus recovering a great number of high-quality bins. Moreover, for each MAG, completeness and contamination are determined using single-copy genes with CheckM (v1.0.12) (Parks et al., 2015). Users can then select MAGs of desired quality for downstream analyses. We select near-complete, high-quality (HQ) genomes as those having ≥90% genome completeness and <5% contamination for downstream analysis.

Taxonomic classification is conducted using GTDB-Tk (v2.1.0) (Chaumeil et al., 2020). GTDB-Tk allows robust taxonomic classification by identifying single-copy marker genes from the Genome Taxonomy Database reference genomes using HMMER (Eddy, 2011). The domain-specific marker gene alignments are then concatenated into multiple sequence alignment and used to construct a reference tree by a maximum likelihood-based phylogenetic inference algorithm. The tool improves the resolution of microbial taxa by classifying MAGs based on their position in the GTDB reference tree, evolutionary divergence, and average nucleotide identity to the reference genomes.

### Functional annotation

Representative cluster centroids of the gene catalogue are annotated with predicted gene functions using two complementary approaches; deep neural networks in DeepFRI (Gligorijevic et al.,2019) and orthology based annotation in eggNOG (evolutionary genealogy of genes: Non-supervised Orthologous Groups) (Huerta-Cepas et al., 2019). eggNOG provides accurate function by inferring orthology relationships and evolutionary history of proteins and is widely used for gene functional annotations. The annotated genes are mapped to various ontologies thus accurately assigning each gene to its function. These ontologies include Gene Ontology (GO), Kyoto Encyclopedia of Genes and Genomes (KEGG) for metabolic pathway analysis, COGs and SMART/Pfam domains for each group.

On the other hand, DeepFRI (Gligorijevic et al., 2021) is a novel deep learning-based function annotation method which applies Graph Convolutional Networks (GCNs) to learn and extract features from protein sequences and structures. The method achieves accurate prediction of GO terms in two stages, (1) the first step utilizes long short term memory (LSTM)-based language model, pre-trained on sequences from Pfam database, to extract features from PDB protein sequences, then (2) GCN architecture learns structure to function relationships using three graph convolutional layers. DeepFRI has the ability to predict protein functions accurately irrespective of sequence homology and provide interpretability of its predictions via saliency maps.

### Integration of gene functions, quantification and metagenome assembled genomes

We have developed a custom Python script for generating mapping between functionally annotated non-redundant genes and metagenome assembled genomes (MAGs). The script takes in a variety of input files: gene catalogue including cluster members, assembled contigs and metagenome assembled genomes (MAGs) (including taxonomic annotations from GTDB and CheckM quality control information).

Considering that the gene catalogue was created by clustering genes into a representative sequence (centroid) using CD-HIT (Fu et al., 2012) at a sequence similarity threshold of 95%, we propagate functions within each gene cluster and further annotate genes within MAGs. This establishes a link between microbial genes and species allowing reconstruction of the functional potential of individual species to determine organisms contributing to various functions. Thus, providing a deeper and more comprehensive insight into identification of species-specific genes.

Quantification of individual gene abundances across a given metagenome is critical for understanding how variation in the functional composition can impact human health as well as in understanding how microbes adapt to various environments. We perform quantifications of each gene and then apply the abundance values to quantify gene functions from DeepFRI and eggNOG. To estimate per-sample gene abundances, the quality-controlled reads are aligned directly against the non-redundant gene catalogue KMA tool (Clausen et al., 2018) with default parameters. The number of reads mapping to each gene is then used as a proxy for its abundance in the sample. For example, the total number of reads mapped to each gene is normalized by the length of the target gene sequence to produce counts in copies per million (CPM) units. The relative abundance is calculated as follows:

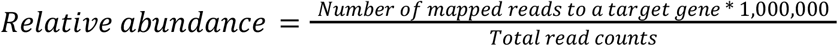

Metagenome assembled genomes (MAGs) include core genes that have specific and specialized function as well as accessory genes that are variably present in the genomes. We use MAGs to construct pan-genomes in reference free manner. To construct a pan-genome of a given species, we recruit near-complete MAGs (≥90% genome completeness and <5%) and species with at least 10 genomes. We then define genes present in ≥90% of MAGs of each species as “core” genes, while genes present in <90% of MAGS are considered as “accessory” genes. This provides additional dimension to analysis of the functional repertoire of core and accessory genomes and gives a glimpse into the functional diversity within species.

## Code availability

The Docker images and scripts used in the workflow are available at https://github.com/microbiome-immunity-project/metagenome_assembly

## Acknowledgements

MM, PS and TK are supported by the Polish National Agency for Academic Exchange grant PPN/PPO/2018/1/00014. TV acknowledges the use of the Broad Institute computing facility.

## Author contributions

MM – performed analyses, contributed analytical tools, wrote the paper

PS – contributed analytical tools, performed analyses

VB – performed analyses, edited the manuscript

VG – contributed analytical tools

CC – contributed analytical tools RB – supervised the project

RJX – contributed analytical tools

TK – conceived the study, supervised the project, wrote the paper

TV – conceived the study, contributed analytical tools, supervised the project, wrote the paper

## Competing interests

All authors declare that there is no competing interest.

## Notes

### Competing Interest Statement

The authors have declared no competing interest.

### Summary of Updates

- we corrected Figures to be of better quality - added an analysis of antibiotics-related functions (main text + Supplement) - corrected the author list - minor text edits

