## Supplementary Information for "Comprehensive functional annotation of metagenomes and microbial genomes using a deep learning-based method"

#### for

### Supplementary figures

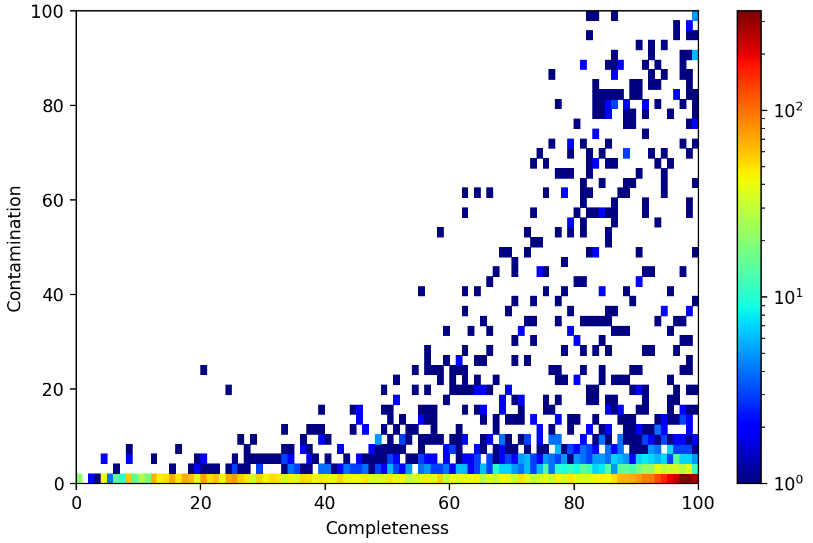


**Supplementary figure 1: CheckM quality estimation of reconstructed genomes.** Completeness and contamination distribution of 7,174 reconstructed MAGs. Genomes with a threshold of completeness ≥90%; contamination <5% were classified as high-quality near-complete genomes.

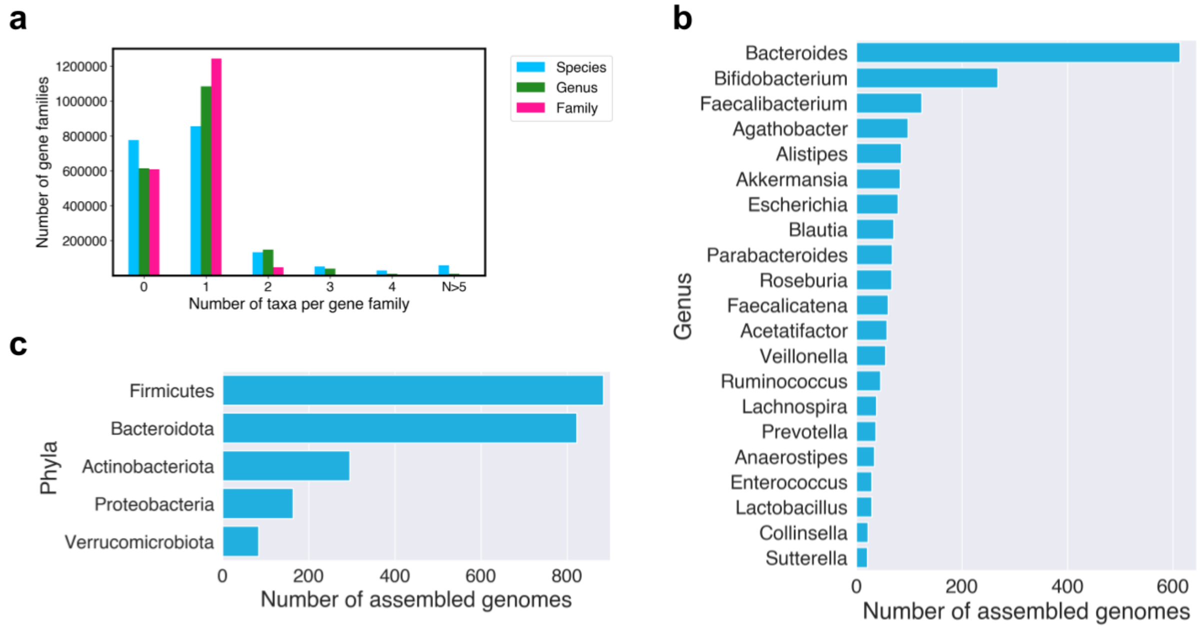


**Supplementary figure 2: GTDB-tk (Chaumeil et al., 2020) taxonomic annotation of gene families and high quality near-complete genomes. (a)** Number of taxonomies per gene cluster. Zero (on x-axis) indicates genes that could not be confidently assigned to any taxonomy**. (b)** GTDB-tk genus assignment of the high quality genomes (top 20 most abundant genus). **(c)** GTDB-tk phyla assignment of the high quality genomes (top 5 most abundant phyla).

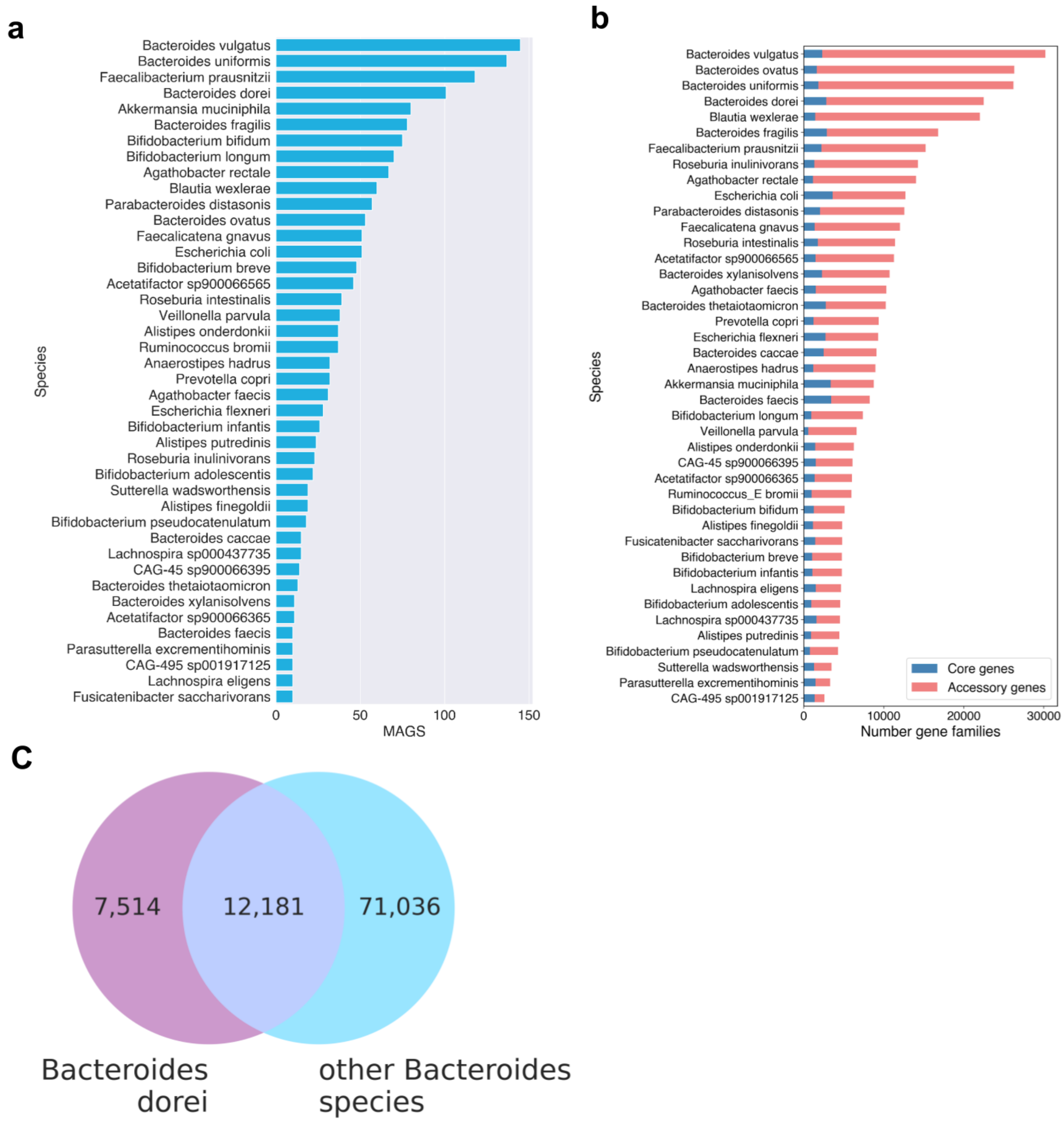


**Supplementary figure 3: Pan-genome statistics. (a)** Species used in pan-genome construction (number of MAGs per species). Entries are ordered according to the number of MAGs. (**b)** Plot of the number of core and accessory genes per species. (**c)** Venn diagram showing unique accessory genes in *Bacteroides dorei* species.

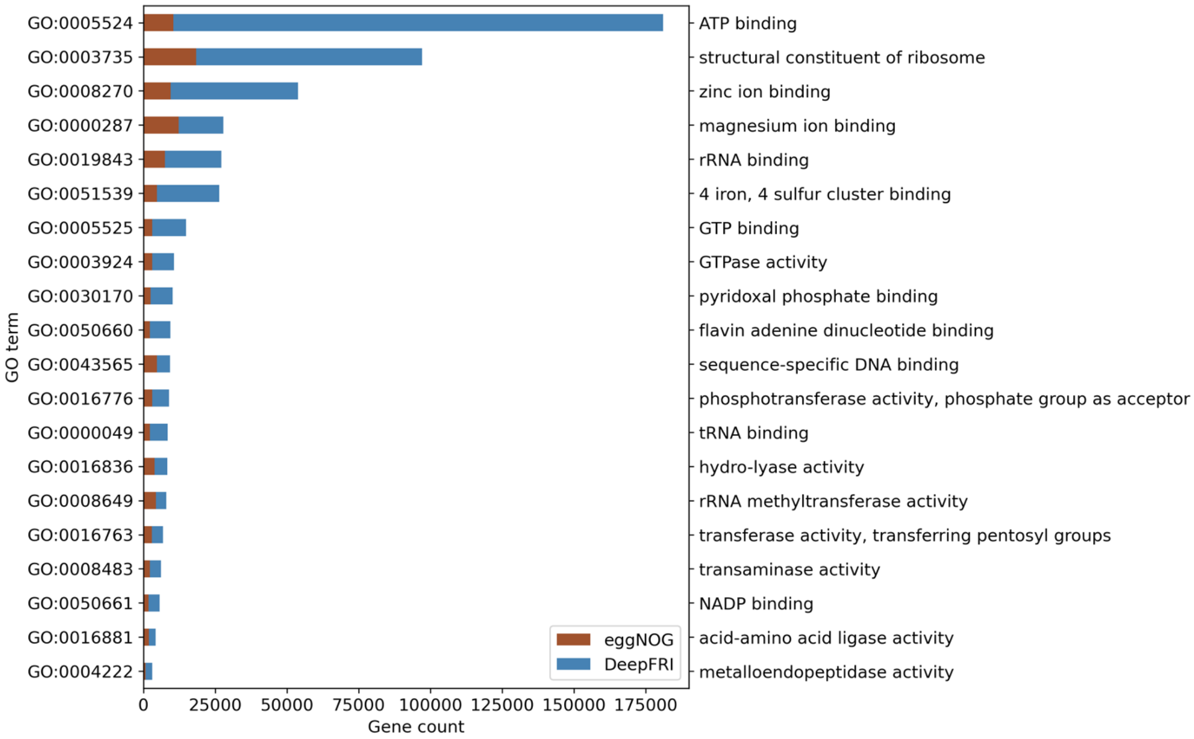


**Supplementary figure 4:** Top twenty most common molecular function GO terms annotated by DeepFRI and eggNOG. See Table S2 for the full list of GO term frequencies.

### Comparisons of DeepFRI (probability threshold 0.5) and eggNOG annotations

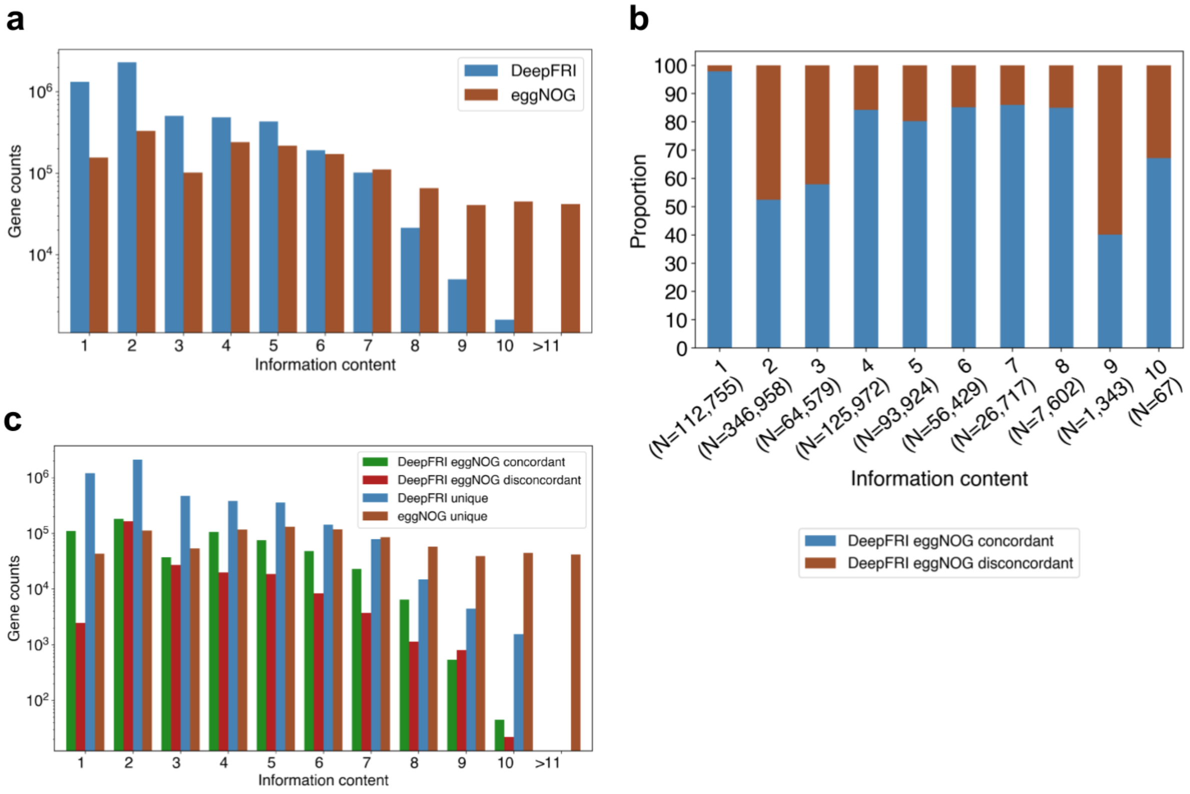


**Supplementary figure 5: Concordance between DeepFRI (threshold 0.5) and eggNOG annotations. (a)** Difference in the annotation information content between eggNOG and DeepFRI **(b)** Proportion of concordant and discordant annotations between DeepFRI and eggNOG per each information content level **(c)** Consensus between DeepFRI and eggNOG annotations.

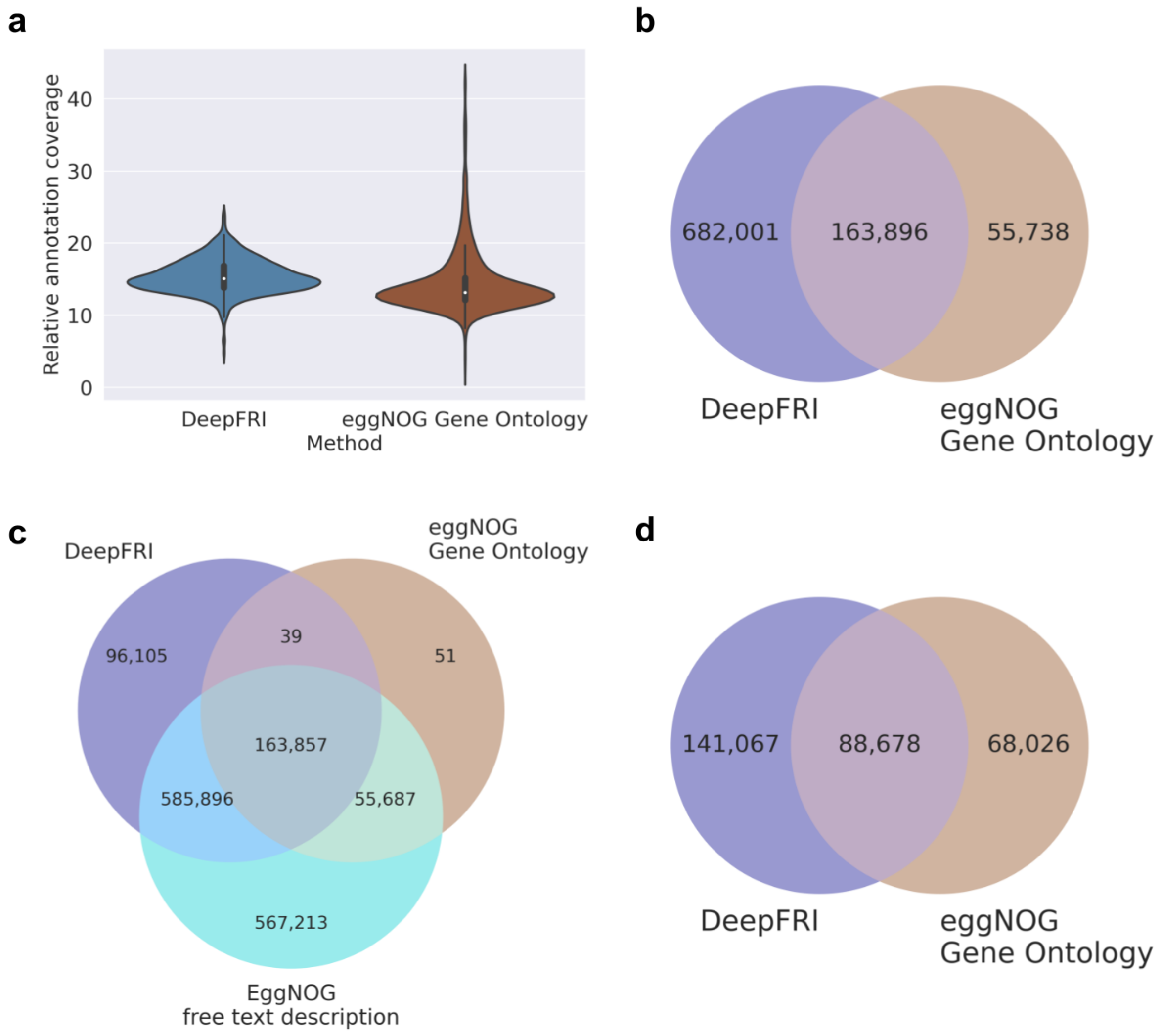


**Supplementary figure 6: Comparisons of functional predictions by DeepFRI (threshold 0.5) and eggNOG. (a)** Proportion of metagenomics gene abundance with functional annotation by DeepFRI and eggNOG (informative gene ontology terms). **(b)** Comparison of gene sets annotated by DeepFRI and eggNOG gene ontology (all GO terms) **(c)** 3-way comparisons of gene sets annotated by DeepFRI, eggNOG gene ontology (all GO terms) and eggNOG free text description (**d)** Venn diagram comparisons of gene sets annotated by DeepFRI and eggNOG using only informative gene ontology terms.

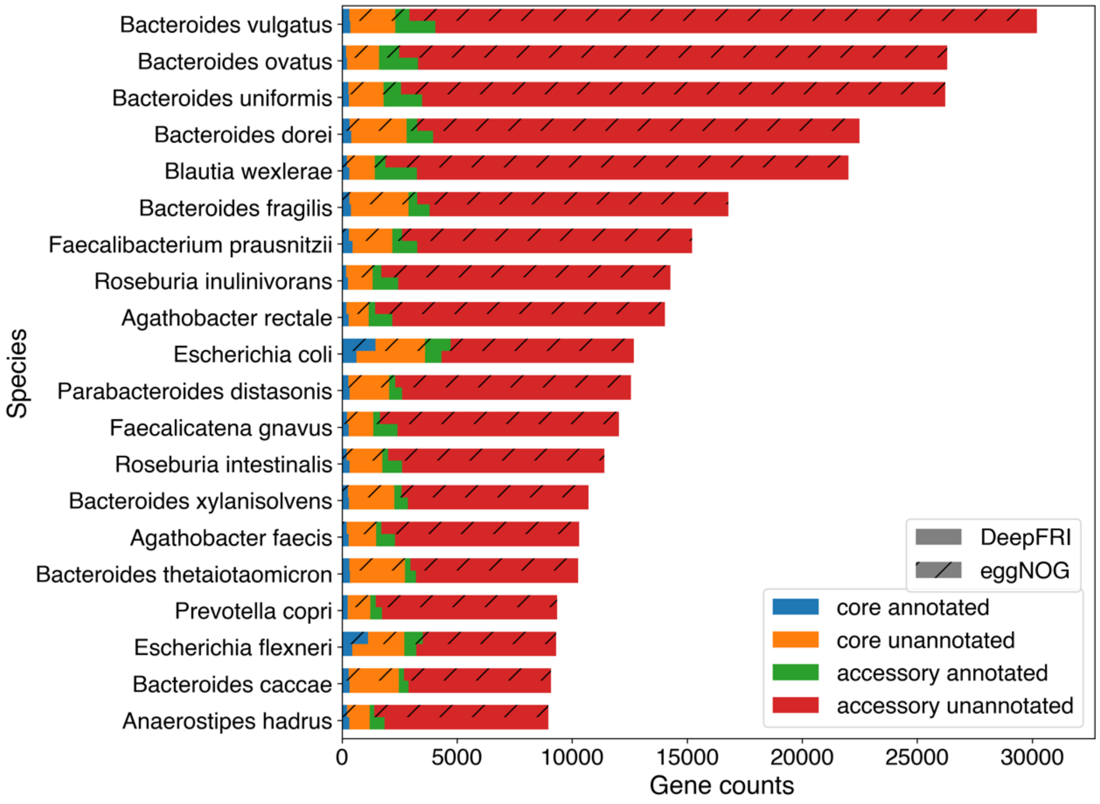


**Supplementary figure 7: Functional annotation of core and accessory genome.** Core and accessory genomes stratified by the functional annotation of genes using eggNOG and DeepFRI, threshold 0.5 (known versus unknown function).

#### Supplementary tables

**Table S1:** Pan-genome size and unique accessory genes (XLSX file).

**Table S2:** List of GO term frequencies (XLSX file).

**Table S3:** List of information content (IC) values for GO terms (XLSX file).

**Table S4a:** List of GO terms related to antibiotics (XLSX file).

**Table S4b:** GO terms associated with antibiotic catabolic and metabolic process

| Method | GO term | Name | No. of genes | No. of species |
|---|---|---|---|---|
| **DeepFRI** | GO:0017001 | antibiotic catabolic process | 32 | 22 |
| | GO:0016999 | antibiotic metabolic process | 12 | 12 |
| **EggNOG** | GO:0017001 | antibiotic catabolic process | 307 | 96 |
| | GO:0016999 | antibiotic metabolic process | 1,635 | 232 |

**Table S5:** Genes annotated by DeepFRI (probability threshold 0.5) and eggNOG gene ontology.

| **Method** | **DeepFRI molecular function** | **eggNOG Molecular function** |
|---|---|---|
| **Annotated genes** | 845,897 (44%) | 219,634 (12%) |
